# Generation of isogenic models of Angelman syndrome and Prader-Willi syndrome in CRISPR/Cas9-engineered human embryonic stem cells

**DOI:** 10.1101/2023.08.30.555563

**Authors:** Rachel B. Gilmore, Dea Gorka, Christopher E. Stoddard, Justin L. Cotney, Stormy J Chamberlain

## Abstract

Angelman Syndrome (AS) and Prader-Willi Syndrome (PWS), two distinct neurodevelopmental disorders, result from loss of expression from imprinted genes in the chromosome 15q11-13 locus most commonly caused by a megabase-scale deletion on either the maternal or paternal allele, respectively. Each occurs at an approximate incidence of 1/15,000 to 1/30,000 live births and has a range of debilitating phenotypes. Patient-derived induced pluripotent stem cells (iPSCs) have been valuable tools to understand human-relevant gene regulation at this locus and have contributed to the development of therapeutic approaches for AS. Nonetheless, gaps remain in our understanding of how these deletions contribute to dysregulation and phenotypes of AS and PWS. Variability across cell lines due to donor differences, reprogramming methods, and genetic background make it challenging to fill these gaps in knowledge without substantially increasing the number of cell lines used in the analyses. Isogenic cell lines that differ only by the genetic mutation causing the disease can ease this burden without requiring such a large number of cell lines. Here, we describe the development of isogenic human embryonic stem cell (hESC) lines modeling the most common genetic subtypes of AS and PWS. These lines allow for a facile interrogation of allele-specific gene regulation at the chromosome 15q11-q13 locus. Additionally, these lines are an important resource to identify and test targeted therapeutic approaches for patients with AS and PWS.

## Introduction

Deletions of the maternal or paternal alleles of chromosome 15q11-q13, respectively, cause Angelman Syndrome (AS [OMIM #105830]) and Prader-Willi Syndrome (PWS [OMIM #176270]). Each occur at an approximate incidence of 1/15,000 to 1/30,000 live births (Burd et al., 1990; Clayton-Smith & Pembrey, 1992; Whittington et al., 2001). Clinical features of AS include seizures, intellectual disability, absent speech, ataxia, and characteristic happy demeanor (Angelman, 1965). Other common features include microcephaly, abnormal EEG, sleep disturbances, hypopigmentation, and strabismus (Williams et al., 1995). AS can be attributed to loss of function of *UBE3A* (Kishino et al., 1997; Matsuura et al., 1997). Clinical features of PWS include neonatal hypotonia and failure-to-thrive during infancy, followed by hyperphagia and obesity; small stature, hands and feet; mild to moderate cognitive deficit and behavioral problems similar to obsessive–compulsive disorder (Cassidy & Driscoll, 2009; Holm et al., 1993; Prader et al., 1956). While PWS is generally thought to be a multigenic disorder, recently described microdeletion cases encompassing just the *SNORD116* cluster highlight its crucial role in PWS pathophysiology (Tan et al., 2020). AS and PWS can be caused by a few different molecular mechanisms, but the most common is a large deletion, affecting ∼70% of patients (Glenn et al., 1997; Kim et al., 2012). The chromosome 15q11-q13 locus harbors intrachromosomal segmental duplications that can misalign during meiosis to generate these “common” large deletions within this chromosomal region (Amos-Landgraf et al., 1999).

Many of the genes in this region are governed by genomic imprinting, a phenomenon in which genes are expressed exclusively from one parental allele, rendering them functionally haploid. Deletion of single copies of the expressed alleles of these imprinted genes cause their full loss of function. Genomic imprinting at chromosome 15q11-q13 is established in the germline via differential methylation at the Prader-Willi Syndrome Imprinting Center (PWS-IC) (Brannan I & Bartolomei, 1999; Nicholls et al., 1998; Shemer et al., 2000). The PWS-IC is methylated on the maternal allele and unmethylated on the paternal allele. This region on the unmethylated paternal allele serves as the canonical promoter for *SNRPN* transcript which is exclusively expressed from the paternally inherited allele. The *SNRPN* transcript is bi-cistronic, encoding for *SNURF* and *SNRPN* (Gray et al., 1999), and also codes a long non-coding RNA (lncRNA), *SNHG14* (reviewed by Ariyanfar & Good, 2022). *SNHG14* can be divided into two units, proximal and distal, based on its expression pattern. The proximal unit is broadly expressed across multiple tissue types and includes *SNURF-SNRPN, SNORD107, SNORD64, SNORD108, IPW, SNORD109A*, and *SNORD116*. The distal unit is exclusively expressed in neural cell types and includes *SNORD115, SNORD109B*, and *UBE3A-ATS* (Cavaillé et al., 2000; Runte et al., 2001). The *UBE3A-ATS* portion of the transcript is responsible for silencing the paternal copy of *UBE3A* (Rougeulle et al., 1998), thus expression of *UBE3A* occurs exclusively from the maternal allele in neurons. Many of the encoded RNAs are of the small nucleolar RNA (snoRNA) class, which are generally thought to be processed by exonucleolytic trimming from the introns of a host gene (Kiss & Filipowicz, 1995). *SNORD116* and *SNORD115* are two clusters of snoRNAs, with 30 individual copies and 48 individual copies respectively. *SNORD116* can be further subdivided into three subgroups: Group I (*SNOG1, SNORD116-1* to *SNORD116-9*), Group II (*SNOG2, SNORD116-10* to *SNORD116-24*), and Group III (*SNOG3, SNORD116-25* to *SNORD116-30*)(Castle et al., 2010; Runte et al., 2001). Protein-coding genes *MKRN3, MAGEL2* and *NDN*, are positioned upstream of the PWS-IC and are exclusively expressed from the paternally inherited allele.

The generation of patient-derived induced pluripotent stem cells (iPSCs) has led to increased understanding of human gene regulation at the chromosome 15q11-q13 locus (Chamberlain et al., 2010; Hsiao et al., 2019; Langouët et al., 2018, 2020). However, variability in the genetic background and epigenetic reprogramming between different iPSC lines make it difficult to study the functional consequences of 15q imprinting disorders in neural cells. Here, we report the generation and characterization of isogenic chromosome 15q11-q13 megabase-scale deletions to model the most common genetic subtypes of AS and PWS. These models were built in the well-characterized and user-friendly H9 human embryonic stem cell (hESC) line to make use of the extensive publicly available data (Dunham et al., 2012; Roadmap Epigenomics Consortium et al., 2015) and robust neuronal differentiation (Itskovitz-Eldor et al., 2000; Schuldiner et al., 2001). The use of isogenic cell lines provides a more rigorous approach to investigate cellular deficits in disease models. These cell lines are well-suited for identifying quantitative molecular and physiological phenotypes, increasing confidence that observed differences between disease and control cells are due to the genetic disorders.

## Results

To generate isogenic models of AS and PWS, we sought to recapitulate deletions frequently present in AS or PWS patients. Examination of deletions deposited in ClinVar and DECIPHER (Firth et al., 2009; Landrum et al., 2014) revealed breakpoint hotspots that coincide with repeats of *GOLGA8* (Figure 1). This repeated sequence described as contributing significantly to substantial instability at this locus (Antonacci et al., 2014; Maggiolini et al., 2019). Therefore we pursued targeting these segmental duplications, similar to the approach used to eliminate the Y-chromosome or trisomic chromosome 21 (Adikusuma et al., 2017; Zuo et al., 2017). We previously used a similar CRISPR/Cas9 strategy, with guide RNAs (gRNAs) targeting *GOLGA8* and other repetitive sequences in chromosome 15, to evict an extra chromosome and generate an isogenic model for Duplication 15q Syndrome (Dup15q, [OMIM #608636]) (Elamin et al., 2023). Others have created 15q13.3 microdeletions leveraging this approach (Tai et al., 2016). Building on the concepts utilized in these previous studies, we began by nucleofecting H9 hESCs with a plasmid encoding CRISPR/Cas9 and a single gRNA targeting *GOLGA8* repeats on chromosome 15q (Figure 1)(Methods). This gRNA is predicted to target multiple sites within chromosome 15q but is not predicted to target elsewhere in the genome. We screened clones surviving transient puromycin selection, which eliminated cells that did not receive the Cas9/gRNA plasmid, for expression of *UBE3A* with a TaqMan-based assay (Methods). As stem cells bi-allelically express *UBE3A*, cell lines harboring AS and PWS related deletions should therefore express approximately half as much *UBE3A* as the parent H9 line (Figure 2A). This screening method provided us with a high-throughput way to screen 126 clones from 4 separate transfections (Supplemental Figure 1, Supplemental Table 1). Three clones with reduced *UBE3A* expression comparable to an Angelman iPSC line (ASdel1-0) were expanded and subject to confirmatory testing. While our initial screen utilized cDNA and relative expression of *UBE3A*, we confirmed deletions by determining the copy number of UBE3A in genomic DNA (gDNA) extracted from our edited clones. We compared edited clones to wild type H9 samples with two UBE3A copies and the ASdel1-0 line with only one UBE3A copy. All three clones were predicted to contain a single UBE3A copy (Methods)(Figure 2B, Supplemental Table 2). To determine the parent-of-origin of the deletions in our three cell lines, we subjected gDNA isolated from them to methylation analysis at the Prader-Willi Syndrome Imprinting Center (PWS-IC, *SNRPN*)(Methods). A wild type cell line will show ∼50% methylation at *SNRPN*, as the paternal allele is unmethylated and the maternal allele is methylated. Previous analysis of patient-derived AS and PWS lines showed that, as expected, PWS deletion lines only have a methylated maternal allele and are ∼100% methylated, and AS deletion lines only have an unmethylated paternal allele lacking methylation at this CpG island (Chamberlain et al., 2010). The methylation analysis indicated that two of the three clones with reduced *UBE3A* expression and copy number exhibited a primarily unmethylated PWS-IC resembling AS (H9Δmat15q_1 and H9Δmat15q_2) and one clone exhibited a primarily methylated PWS-IC resembling PWS (H9Δpat15q) (Figure 2C)(Supplemental Table 3). To determine the approximate size of the deletion, clones were further characterized by a CytoSNP analysis. This analysis revealed an ∼5.8Mb deletion in the H9Δmat15q_1 line, an ∼8Mb deletion in the H9Δmat15q_2 line, and two deletions totaling ∼7Mb in the H9Δpat15q line (Supplemental Figure 2). The coordinates of the deletions returned from CytoSNP analysis were displayed as bedtracks in the UCSC genome browser to visually display the size of the deletions and which genes may be impacted (Figure 2D). These deletions coincided well with those observed in patients (Supplemental Figure 3), further supporting *GOLGA8*-driven instability as genetic mechanism leading to AS and PWS.

**Figure 1.**
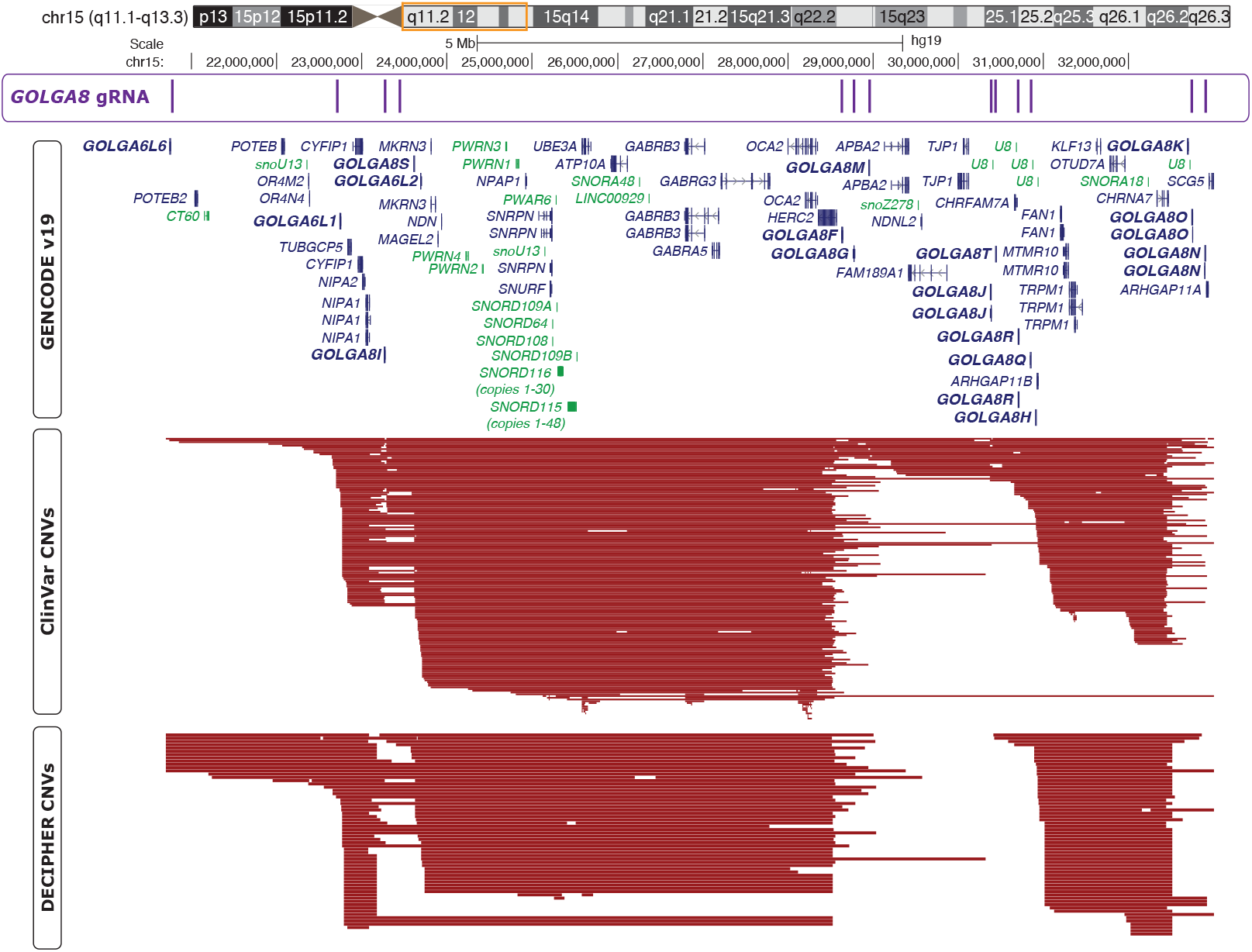
UCSC Genome Browser shot of the chromosome 15q locus. Chromosome ideogram with orange box around region displayed below. First track displays *GOLGA8* CRISPR gRNA binding sites. Second track displays GENCODEv19 gene annotations; genes in dark blue are protein coding, genes in green are non-coding, arrows indicate direction of gene transcription. Some isoforms for genes are removed for clarity. GOLGA genes are shown in bold. Third track displays ClinVar copy number variants (CNVs); only deletions with pathogenic or likely pathogenic annotations are shown. Fourth track shows DECIPHER CNVs, only deletions with pathogenic or likely pathogenic annotations are shown.

**Figure 2.**
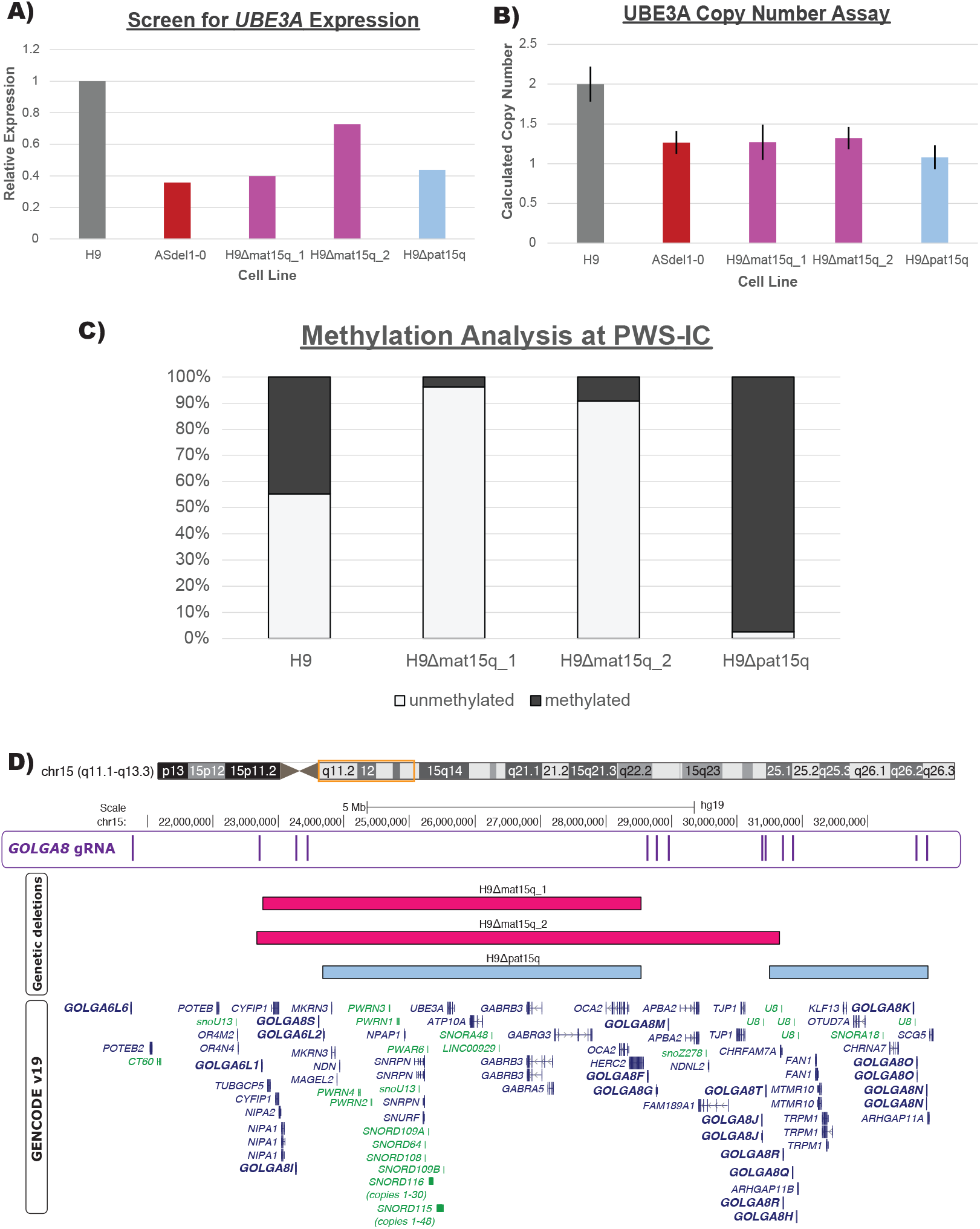
**A)** Bar plot of relative *UBE3A* expression of edited clones compared to wild type (H9) ESCs and Angelman Syndrome (ASdel1-0) iPSCs. **B)** Bar plot of calculated copy number of edited clones as determined by CopyCallerv1.2 software. Wild type (H9) was used as the calibrator sample and set to 2 copies. Error bars represent calculated copy number range as determined by CopyCallerv2.1 software. **C)** Stacked bar plot of methylation analysis at the PWS-IC/*SNRPN* locus. Light gray bar represents unmethylated DNA, dark gray bar represents methylated DNA. **D)** UCSC Genome Browser shot of the chromosome 15q locus. First track displays *GOLGA8* CRISPR gRNA binding sites. Second, third, and fourth tracks show deletion contained within each cell line, as determined by CytoSNP. Fifth track displays GENCODEv19 gene annotations; genes in dark blue are protein coding, genes in green are non-coding, arrows indicate direction of gene transcription. Some isoforms for genes are removed for clarity. GOLGA genes are shown in bold.

**Figure 3.**
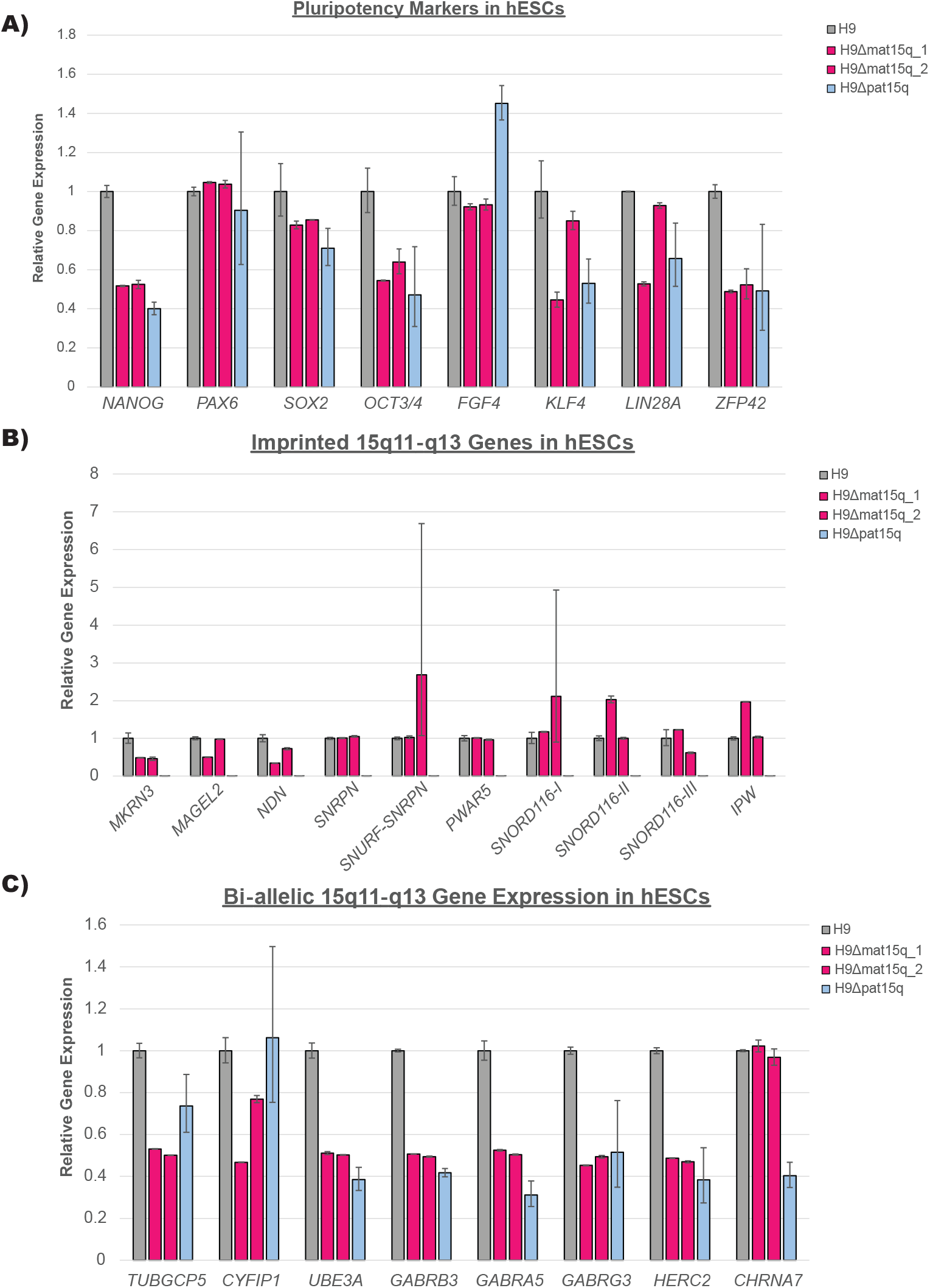
qPCR analysis of **A)** pluripotency markers, **B)** imprinted genes in the 15q locus, and **C)** bi-allelically expressed genes in the 15q locus in edited clones as ESCs (n = 2 biological replicates). RNA expression is presented relative to the parental wild type H9 ESC line. Error bars represent standard error of the mean ΔCt.

We next sought to determine whether these isogenic lines still showed pluripotent potential and to characterize their gene expression profile in the chromosome 15q locus in ESCs (Methods)(Supplemental Table 4 & 5). None of the cell lines showed distinct differences in pluripotency markers compared to the parental H9 line (Figure 3A). In lines with maternal deletions, imprinted 15q genes showed similar expression to the parental H9 line as expected (Figure 3B). In the paternal deletion line, imprinted 15q genes that are expressed exclusively from the paternal allele showed very little expression compared to the parental H9 line. This is expected, as these imprinted genes are silenced on the intact maternal allele (Figure 3B). This data supported our characterization of which allele was deleted in each cell line. The bi-allelically expressed 15q genes within the deletion breakpoints showed approximately half of the expression of the parental H9 line (Figure 3C). As differentiation into the neuronal lineage is important for studying these disorders and is crucial for verifying the imprinting status of *UBE3A* in neurons, we selected the H9Δmat15q_1 line for neuronal differentiation and further gene expression characterization due to the more clinically-relevant size of the deletion (Methods)(Supplemental Table 4 & 6). The ESCs successfully differentiated into neurons (Figure 4A). As expected, imprinted 15q gene expression was comparable to that of the parental H9 line, except *UBE3A*, which was drastically reduced (Figure 4B). We considered this proof-of-concept that these isogenic cell line models are capable of imprinting *UBE3A* following neuronal differentiation. Bi-allelic 15q gene expression maintained a similar expression profile to that observed in ESCs, with genes contained within the deletion showing approximately half expression compared to the parental H9 line (Figure 4C). Pluripotency markers, except for *PAX6*, show decreased expression in neurons compared to wild type ESCs (Supplemental Figure 4). Given that these cell lines readily differentiate into neurons, we anticipate they will provide great utility for understanding the specific role these deletions play in neurodevelopmental phenotypes of AS and PWS.

**Figure 4.**
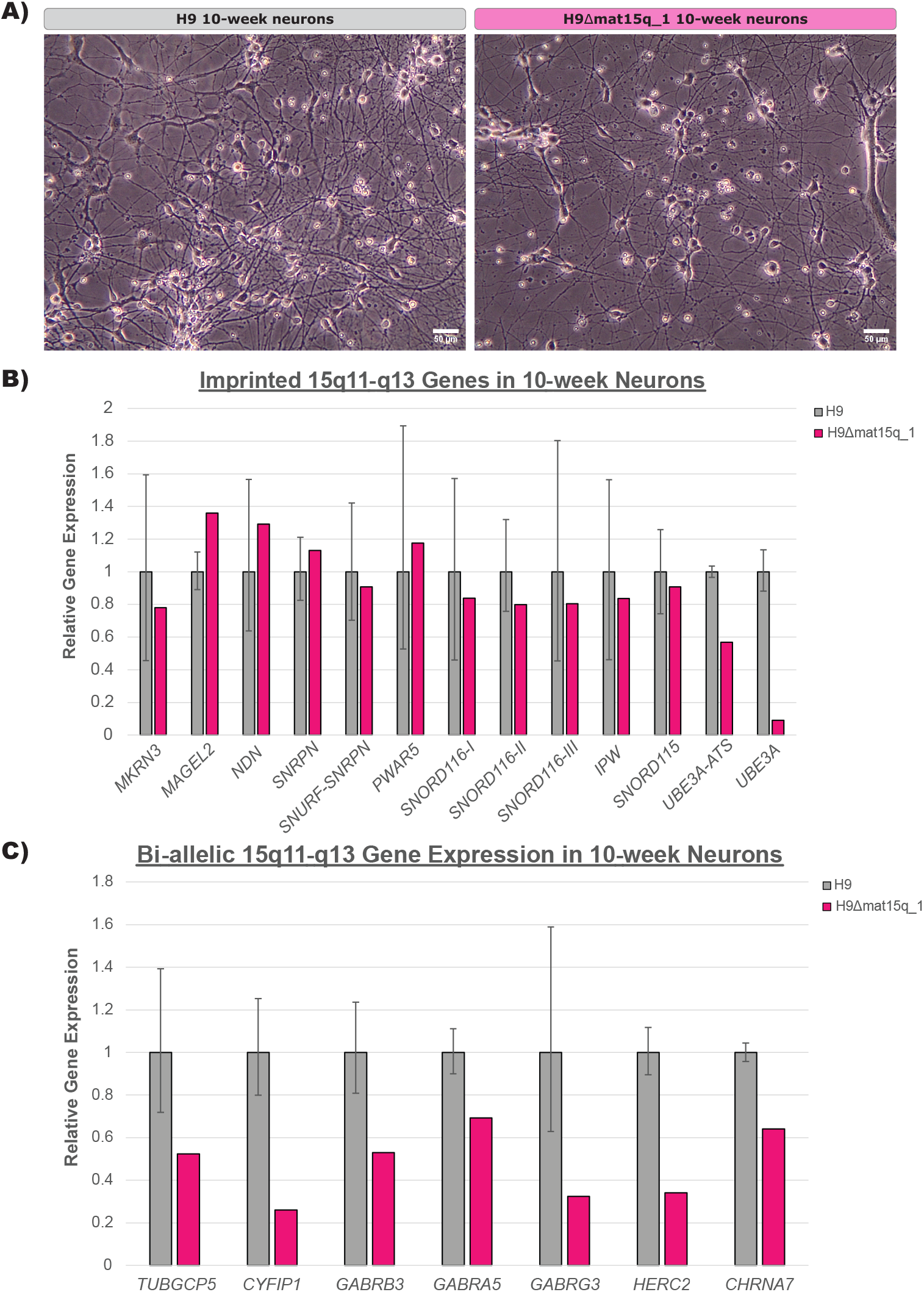
**A)** Representative brightfield images of wild type (H9) and maternal deletion line neurons at 20X magnification. Scale bar equals 50um. **B&C)** qPCR analysis of **B)** imprinted genes and **C)** bi-allelically expressed genes in the 15q locus in edited clones as mature 10-week neurons (n = 1-2 biological replicates). RNA expression is presented relative to the parental H9 line. Error bars represent standard error of the mean ΔCt.

## Discussion

While the genetic perturbations contributing to AS and PWS have been known for many years, the effect those anomalies have on the chromosome 15q locus and the genome as a whole remains unclear. Mouse models have provided key understandings about the facets of gene regulations conserved between the two species, but disease features such as the neuron-specific regulation of *UBE3A* imprinted expression and protein targets of UBE3A appear to be unique to humans (Cavaillé et al., 2000; Dindot et al., 2023; Landers et al., 2004; Yamasaki et al., 2003). Available iPSC models have been powerful tools to study these disorders as well. However, comparison of quantitative molecular phenotypes via multiomics approaches and functional studies of iPSC-derived neurons have been hampered by the variability between iPSC lines. Here we have described the first isogenic cell line pairs modeling the most common genetic subtypes of AS and PWS. We reasoned that we may be able to mimic megabase-scale deletions found in patients by targeting chromosome 15q-specific *GOLGA8* repeats in a well-characterized H9 ESC line. We took advantage of the bi-allelic expression of *UBE3A*, a gene included in the deleted region, in hESCs to rapidly screen for edited clones with reduced expression (Figure 2A, Supplemental Figure 1). Our rationale was that if either the maternal or paternal chromosome 15q allele was deleted in this region, we would observe approximately half the *UBE3A* expression compared to a wild type control. We subjected the clones with reduced *UBE3A* expression to a more rigorous confirmatory testing utilizing a copy number assay to determine the number of *UBE3A* copies present in gDNA extracted from each clone (Figure 2B).

As the parent-of-origin of the deletion matters, we utilized differential methylation at the PWS-IC/*SNRPN* to determine which allele was deleted. The differential methylation at this site has been characterized previously in patient-derived iPSCs (Chamberlain et al., 2010). We hypothesized our isogenic models would have comparable methylation signatures to iPSC models if they harbored similar deletions, which was what we observed (Figure 2C). Confident we created deletions on either the paternal or maternal allele, we employed a CytoSNP array to determine the approximate size of the deletion. This assay leveraged microarray technology to detect copy-neutral loss of heterozygosity (LOH), absence of heterozygosity (AOH), and copy number variation (CNV) via gains or losses. While off-target editing is sometimes a concern with CRIPSR/Cas9 editing, CytoSNP analysis did not detect any copy number changes or structural rearrangements aside from the deletions on chromosome 15. Any copy number changes outside of chromosome 15 in the edited hESCs were also present in the parental H9 cell line previously characterized by the lab, supporting the isogenic nature of these cell lines (Supplemental Figure 2).

Having determined the parent-of-origin and approximate size of the deletions, we wanted to ensure these edited cell lines were still pluripotent. While the expression profile of the pluripotency markers in edited cell lines did not match exactly to the wild type controls (Figure 3A), this could be caused by differences in the quality of the cultures at the time of collection. The pluripotent potential of the edited cell lines was supported by their ability to differentiate successfully into a neuronal lineage (Figure 4A). We also wanted to determine gene expression more accurately within the chromosome 15q locus. All imprinted genes measured in the array (Figure 3B) were contained within the deletions created in each of the three edited lines. We would expect the gene expression in maternal deletion lines to not vary greatly from the parental wild type line, as the imprinted and unexpressed copy of each gene was deleted. However, we noticed decreased expression of *MRKN3, MAGEL2*, and *NDN* in these lines, which normalized to near wild type levels in neurons (Figure 4B). Further study of these cell lines at a chromatin level may reveal whether this peculiar gene expression pattern frequently occurs in iPSCs/hESCs with maternal 15q deletions or whether it is unique to these engineered cell lines. In contrast, we would anticipate little to no expression of the paternally expressed imprinted genes in the paternal deletion line, as the expressed copy of each gene was deleted, which was exactly what we observed (Figure 3B). As *SNORD115* and *UBE3A-ATS* are not expressed in ESCs (Supplemental Table 5), these genes were only included in the analysis of neurons (Figure 4B).

The bi-allelically expressed genes, *TUBGCP5* and *CYFIP1* were only deleted in the two maternal deletion lines, and showed expression reduced by approximately half in ESCs (Figure 3C) and neurons (Figure 4C) as expected. In the paternal deletion line, these two genes showed expression levels similar to the parental H9 hESC line (Figure 3C). *GABRB3, GABRA5, GABRG3* were included in all three edited lines and showed expression reduced by approximately half in ESCs (Figure 3C) and neurons (Figure 4C) as expected. While *HERC2* was only entirely deleted in the H9Δmat15q_2 line, the other two edited lines showed similar reduction in expression with deletion of exons 5-93 (Figure 3C). We would anticipate whatever protein product produced from the remaining portion, if any, to be non-functional. *CHRNA7* was only contained within the paternal deletion line, and therefore only showed reduced expression in that line (Figure 3C) as expected. However, we also noted a slight decrease in expression of *CHRNA7* in neurons generated from the maternal deletion line (Figure 4C), which may suggest differential regulation of this gene in neurons compared to ESCs. While *UBE3A* is bi-allelically expressed in ESCs, which we exploited for rapid screening of clones, the *UBE3A* copy on the paternal allele undergoes silencing in neurons (Cavaillé et al., 2000; Chamberlain et al., 2010; Rougeulle et al., 1998; Runte et al., 2001). We showed that one of our maternal deletion lines successfully differentiated into neurons and showed evidence of *UBE3A* imprinting (Figure 4A&B). Support of the successful neuron differentiation was typical neuron morphology and the decrease in expression of pluripotency markers (Figure 4A, Supplemental Figure 4). *PAX6* showed increased expression in both wild type neurons and maternal deletion neurons, likely because it has been shown to play a role in neuroectoderm development (Zhang et al., 2010). Further functional studies of these hESC-derived neurons could determine if they display similar deficits to those found in iPSC-derived neurons (Fink et al., 2017).

These isogenic cell lines provide a powerful resource to carefully discern cellular and molecular phenotypes between disease and wild type states for these large chromosomal deletions. We posit that the use of these isogenic pairs will lead to more robust and reproducible results, especially when combined with additional isogenic pairs and/or patient-derived iPSC lines. Additionally, these data may open the door for the discovery of novel, more specific therapeutic approaches for AS and PWS patients. Lastly, this work adds to the current literature supporting the utility of CRISPR/Cas9 editing to eliminate large regions of the genome, which could potentially be applied to other disorders with large copy number variants.

## Materials and Methods

### hESC culture

hESC were maintained on mitotically inactivated mouse embryonic fibroblasts (MEFs) in feeding media which consists of sterile-filtered DMEM/F12 media (Gibco, # 11330032) supplemented with 20% Knock Out Serum Replacement (Gibco, #), 1X MEM Non-essential amino acids (Gibco, #11140050), 1mM L-glutamine (Gibco, #25030081) with 0.14% β-mercaptoethanol, and 8ng/mL bFGF (Gibco, #PHG0023). A humidified incubator with 5% CO_2_ was used to maintain the cells at 37°C. Stem cells were manually passaged by cutting and pasting colonies every 6 or 7 days using a 28-gauge needle. Stem cell media was replaced daily.

### Genome editing of hESCs

H9 ESCs were engineered with a megabase-scale deletion on either the paternal or maternal chromosome 15q allele. A similar editing and screening strategy has been previously described (Elamin et al., 2023).

#### Preparation

A guide RNA targeting *GOLGA8* was designed using available guide RNA design tools (*GOLGA8* gRNA: CTGGGTGTGAGGGCACGTGG). The guide was cloned into the pSpCas9(BB)-2A-Puro (PX459) V2.0 plasmid (Addgene, #62988) via restriction digestion and ligation. Two days prior to planned genome editing, a 100mm dish of mitotically inactivated DR4 MEFs was prepared. A ∼60-75% confluent well of hESCs was treated 24 hours prior to planned genome editing with 10µM ROCK inhibitor, Y-27632 2HCl (Tocris #1254).

#### Nucleofection

The day of editing, one ∼75% confluent well of a 6-well plate of hESCs was treated with Accutase (Millipore Sigma, #SCR005) to release the cells from the plate. The cell suspension was singularized by pipetting and then pelleted. The media was removed from the cell pellet, and cells were resuspended according to the protocol provided for the P3 Primary Cell 4D-Nucleofector Kit (Lonza, V4XP-3024). Briefly, a mixture of 82µL nucleofector solution, 18µL nucleofection supplement, and ∼5 µg of CRISPR plasmid was added to the pellet. The pellet was resuspended in the solution by pipetting gently three times using a P200 pipet. The cell suspension was transferred to the nucleofection cuvette and nucleofection was performed on the 4D-Nucleofector (Lonza) on the program for hESC, P3 primary cell protocol. Atier nucleofection, hESC suspension was immediately transferred to the 100mm dish plated with DR4 MEFs containing hESC feeding media supplemented with 10µM ROCK inhibitor using the transfer pipet included in the kit.

#### Selection

Feeding media was changed 24 hours following transfection (Day 1 post-transfection) and supplemented with 0.5-1 ng/μL puromycin and 10µM ROCK inhibitor. This selection was continued for 48 hours total to select cells transiently expressing the vector containing the gRNA and Cas9 protein. On Day 2, the media was changed and supplemented with fresh 0.5-1 ng/μL puromycin and 10µM ROCK inhibitor. On Day 3, the media was changed and supplemented with fresh 10µM ROCK inhibitor. Subsequent media changes occurred every other day, supplemented with fresh 10µM ROCK inhibitor. Once small colonies became visible, media changes occurred daily with fresh media alone. After ∼2 weeks, each colony was manually passaged into its own well of a 24-well plate coated with MEFs via cutting and pasting. Feeding media in the 24-well plate was supplemented with 10µM ROCK inhibitor to encourage cell attachment. 48 hours after passaging cells, the feeding media was changed. Approximately 4 days after passaging to a 24-well plate, a few colonies from each well were isolated into PCR tube strips and pelleted for screening.

#### Screening

The TaqMan® Gene Expression Cells-to-CT™ Kit (Invitrogen™, #4399002) was used to screen clones following manufacturer’s protocol. Briefly, media was removed from cell pellets in PCR tube strips and diluted DNase I lysis solution was added. A reverse transcription (RT) reaction was performed. Finally, a real-time PCR was run utilizing TaqMan™ Assays to measure expression of *UBE3A* (Hs00166580_m1) (ThermoFisher, #4331182) versus *GAPDH* (ThermoFisher, #4352934E) in technical duplicates or triplicates at a total reaction volume of 20μL. Clones that were found to have ∼50% reduction in *UBE3A* compared to wild type controls were further expanded and subjected to confirmatory testing. A previously described iPSC line derived from an AS patient (ASdel1-0)(Chamberlain et al., 2010) was included in the assay as a control line for half *UBE3A* expression.

#### Confirmatory Testing

After manually passing clones for expansion and verifying attachment of colonies in new wells, the remainder of cell colonies were scraped from old wells and pelleted in microcentrifuge tubes. Genomic DNA (gDNA) was extracted from clones using a homemade lysis buffer containing 1% sodium dodecyl sulfate (SDS), 75mM NaCl, 25mM EDTA, and 200μg/mL Proteinase K in UltraPure™ DNase/RNase-Free Distilled Water (ThermoFisher, #10977015). Briefly, media was removed from each cell pellet and 250uL of the lysis buffer was added. Tubes were incubated at 60°C overnight. The following day, 85μL of warm 6M (supersaturated) NaCl was added, followed by the addition of 335μL of chloroform. The tubes were capped and then inverted for approximately one minute. Tubes were centrifuged at 9,000 rcf for 10 minutes at room temperature. The top aqueous layer (∼335μL) was removed and transferred to a new tube to which an equal amount of 100% isopropanol was added. The tubes were capped and mixed thoroughly by inversion. Tubes were incubated at -20°C for ∼10 minutes. Next, the tubes were centrifuged at max speed (∼18,000 rcf) for 20 minutes at 4°C. The supernatant was removed, and the pellet was washed with ∼600μL of 70% ethanol. Ethanol was removed carefully from the pellet and the tubes were left open so the remainder of the ethanol could evaporate before the pellets were resuspended in 30μL of 10mM Tris (pH 8). TaqMan™ Copy Number Assays comparing UBE3A (Hs01665678_cn)(ThermoFisher, #4400291) to the TaqMan Copy Number Reference Assay for human RNase P (ThermoFisher, #4403326) was performed following manufacturer’s protocol (Applied Biosystems™, Publication Number #4397425) to confirm which clones had lost a copy of *UBE3A*. The wild type H9 line was used as the calibrator sample with two copies of *UBE3A*. The same AS1-0 iPSC line used as a control in our screening assay was also included as a control in this assay as it only has one copy of *UBE3A*. Analysis was conducted using the CopyCaller (v2.1) software (Applied Biosystems®). Further confirmation of which allele was deleted in clones with only one *UBE3A* copy was performed by utilizing the EpiTect Methyl II DNA Restriction Kit (QIAGEN, #335452) to measure methylation at the PWS-IC (*SNRPN*) following manufacturer’s protocols. The approximate size of the deletions was determined by CytoSNP array (Illumina, CytoSNP-850K v1.2) through the University of Connecticut Chromosome Core. Clones with confirmed deletions were expanded, banked down, and subsequently characterized via gene expression arrays in stem cells and neurons.

### Neuronal differentiation and maintenance

Neuronal differentiation was performed according to established monolayer differentiation protocols with minor modifications (Chen et al., 2016; Noelle D. Germain et al., 2014; Noélle D. Germain et al., 2013; Sirois et al., 2020). Approximately 1-3 days after passaging hESCs, neuronal differentiation was started (Day 0) by switching feeding media to N2B27 neural induction media supplemented with 500 ng/μL noggin (R&D Systems, #3344-NG). N2B27 neural induction media consisted of Neurobasal™ Medium (Gibco, #21103049), 1X serum free B-27™ Supplement (Gibco, #17504044), 1% N2 supplement, 1% insulin-transferrin-selenium (Gibco, #51300044), 2mM L-glutamine (Gibco, #25030081), and 1% penicillin-streptomycin (Gibco, #15140122). The media was changed every other day for 10 days and supplemented with fresh 500 ng/μL noggin on Days 2, 4, 6, and 8. Between Days 14-17, neuronal rosettes were passaged as small clusters in either a 1:1 or 1:2 ratio using the StemPro™ EZPassage™ Disposable Stem Cell Passaging Tool (Gibco, #23181010) or via hand-picking. Rosettes were plated on poly-D-lysine-(PDL-)(Millipore Sigma, #P0899) and laminin-coated (Gibco, # 23017015) 6-well plates. Fifty percent media replacement was carried out every other day until neural progenitor cells (NPCs) were dense enough for replating. At ∼3 weeks, Accutase was used to release the cells from the plate. The cell suspension was singularized by pipetting and then pelleted. The media was removed from the cell pellet, NPCs were resuspended, and replated at a high density onto poly-D-lysine/laminin-coated 6-well plates into N2B27 media containing 10µM ROCK inhibitor. Fitiy percent media replacement was carried out every other day. After approximately five weeks of neural differentiation NPCs were dissociated again using Accutase, counted using a hemocytometer, and plated on PDL/laminin-coated 6-well plates with at a density of 150,000-300,000 cells/well in neural differentiation medium (NDM). NDM consisted of Neurobasal™ Medium, 1X serum free B-27™ Supplement, 1X MEM Non-essential amino acids, 2mM L-glutamine, 10ng/mL brain-derived neurotrophic factor (BDNF)(Peprotech, #450-02), and 10ng/mL glial-derived neurotrophic factor (GDNF)(Peprotech, #450-10), 200µM ascorbic acid (Millipore Sigma, #A4544), and 1µM adenosine 3’,5’-cyclic monophosphate (cAMP)(Millipore Sigma, #A9501). To aid in cell attachment, 10µM ROCK inhibitor was added to the NDM during initial plating. Cells were maintained with no antibiotics on NDM, and 50% media replacement was carried out twice per week. Gene expression assays were conducted on neuronal cultures that were at least 10 weeks old.

### Gene Expression Arrays

#### RNA Extraction

Confluent hESCs or mature hESC-derived neurons were collected using RNA-Bee (AMSBIO, #CS-501B) or RNA STAT-60 (AMSBIO, #CS-502) reagent and total RNA was isolated following manufacturer’s protocol with minor adaptations. Briefly, samples were lysed directly in 6-well plates by removing media and adding 1mL of lysis reagent per well. Samples were homogenized by pipetting several times using a P1000 pipet. Samples were incubated for 5 minutes at room temperature. Samples were transferred to microcentrifuge tubes and 200µL of chloroform was added to each. Tubes were capped, shaken vigorously for 30 seconds, and stored on ice for 5 minutes. Samples were centrifuged at 12,000 rcf for 15 minutes at 4°C. The top aqueous layer was removed and transferred to a new tube to which 500µL 100% isopropanol and 2µL of 5mg/mL glycogen (ThermoFisher, #AM9510) was added. Tubes were capped, inverted gently to mix, and stored on ice for 30 minutes for hESCs or up to an hour for hESC-derived neurons. After incubation, samples were centrifuged at 4°C at max speed (∼18,000 rcf) for 30 minutes for hESCs or up to 45 mintues for hESC-derived neurons. The supernatant was removed and the RNA pellet was washed with 1mL of 75% ethanol by shaking the capped tube to dislodge the pellet. Samples were centrifuged at 7,500 rcf for 5 minutes at 4°C. For hESCs, the wash and spin was repeated to improve the purity of the RNA. The extra wash was omitted for neurons to prevent losing RNA yield. Ethanol was removed from pellets and the pellets were air dried briefly at room temperature. When the edges of the pellet became slightly opaque, the RNA pellet was dissolved in a minimum of 11µL of UltraPure™ DNase/RNase-Free Distilled Water. For hESCs, biological duplicates were used. For neurons, 1-2 differentiations were performed. All samples were analyzed in technical duplicates.

#### RT-qPCR

Samples were treated with DNase I (Invitrogen™, #18068015). The High-capacity cDNA Reverse Transcription Kit (Applied Biosystems™, #4368814) was used to generate cDNA from DNase-treated RNA following manufacturer’s protocol. Custom TaqMan Gene Expression Assay Plates (ThermoFisher) were used with TaqMan™ Gene Expression Master Mix (Applied Biosystems™, #4369016) to measure gene expression following manufacturer’s protocol. The list of TaqMan probe RT-qPCR assays used in the study are provided in Supplemental Table 4.

## Data Analysis

For qPCR, the mean Ct value of technical replicates for each gene were normalized to the mean Ct value of technical replicates for housekeeping gene *GAPDH*. Relative expression was quantified as 2^-ΔΔCt^ relative to a wild type H9 sample. Data are presented as the mean relative expression, plus or minus the standard error of the mean ΔCt when applicable.

## Resource availability

Cell lines are available upon reasonable request and after completion of Material Transfer Agreements through the University of Connecticut Cell and Genome Engineering Core. UCSC browser session used for generation of Figure 1, Figure 2D, and Supplemental Figure 3 is publicly available at https://genome.ucsc.edu/s/rbgilmore/Megabase_Deletion_Lines_Figure_ClinVar.

## Author Contributions

S.J.C. conceived of and designed the study. R.B.G. and C.S. carried out genome editing to generate the isogenic cell lines. R.B.G. and D.G. conducted experiments, analyzed data, and wrote the manuscript. All authors were involved in the preparation and review of the manuscript and approved the final submitted version.

## Acknowledgements

This work was supported by National Institutes of Health (NIH) grant R01 HD099975 (J.L.C. and S.J.C.). This study makes use of data generated by the DECIPHER community. A full list of centres who contributed to the generation of the data is available from http://deciphergenomics.org/about/stats and via email from. DECIPHER is hosted by EMBL-EBI and funding for the DECIPHER project was provided by the Wellcome Trust [grant number WT223718/Z/21/Z]. We thank Yaling Liu form the University of Connecticut Cell and Genome Engineering Core for her help with expansion and storage of stem cell lines used in this study.

