## Supplemental Figures for "Generation of isogenic models of Angelman syndrome and Prader-Willi syndrome in CRISPR/Cas9-engineered human embryonic stem cells"

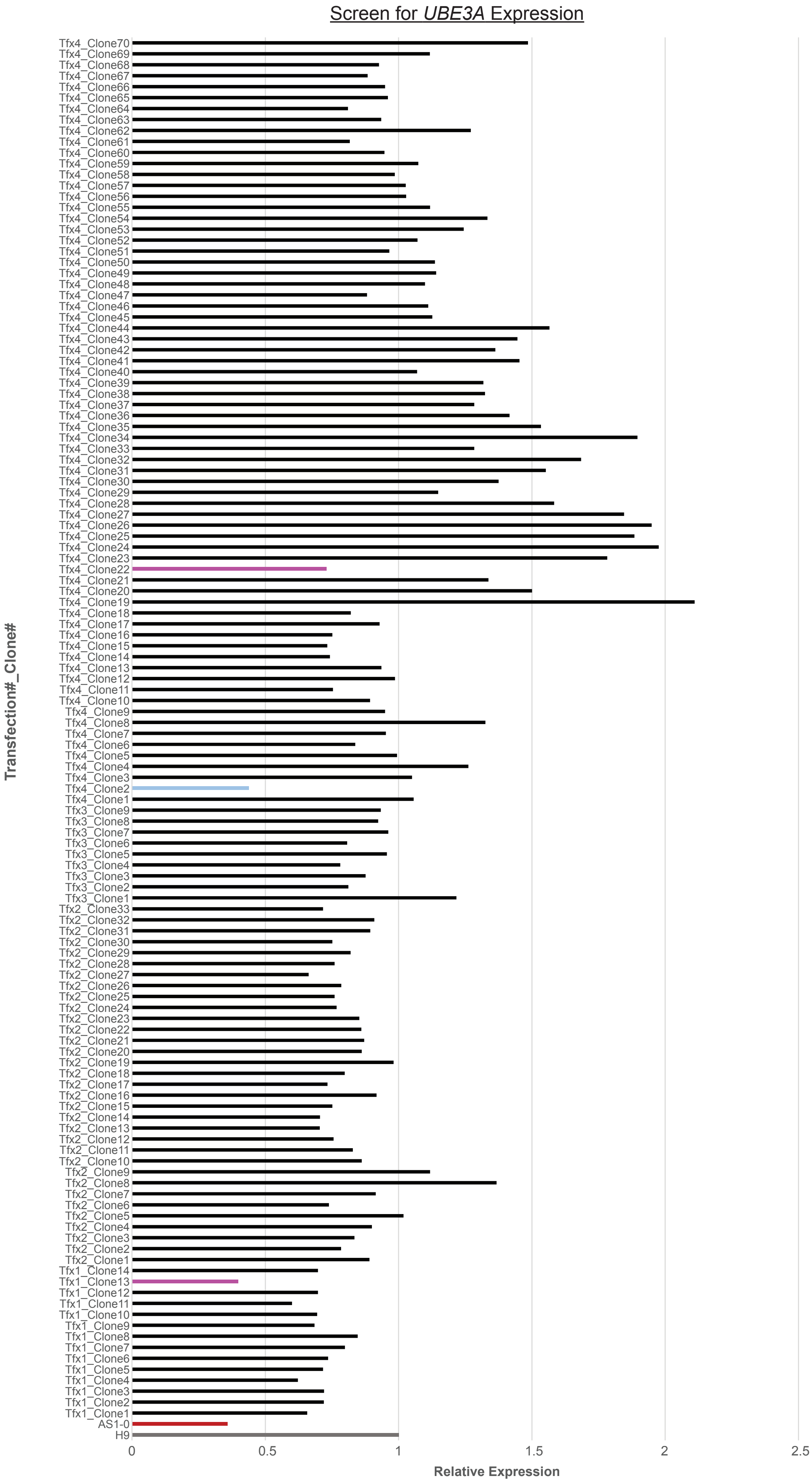


**Supplemental Figure 1.** Compilation of screening data from *UBE3A* expression from all clones from all transfections Relative expression of *UBE3A* is set to a relative value of 1 for the wild type parental H9 line. H9Δmat15q_1 is designated as Tfx1_Clone13, H9Δmat15q_2 is designated as Tfx4_Clone22, H9Δpat15q is designated as Tfx4_Clone2.

UCONN UNIVERSITY OF CONNECTICUT — University of Connecticut Chromosome Core — Chromosome Core

**Case Report**

| | |
|---|---|
| **Sample ID:** | **CC20-11** |
| **Sample Name:** | **H9 AG #22** |
| **Sample arrival date:** | July 11, 2020 |
| **Experiment date:** | August 26, 2020 |
| **Report date:** | April 29, 2020 |
| **Microarray type:** | Illumina CytoSNP-850K v1.2 |
| **Microarray Barcode:** | 204556110015 |
| SNP manifest file: | CytoSNP-850Kv1-2_NS550_B3.bpm |
| Annotation DB: | BG_Annotation_Ens74_20180801.db |
| SNP cluster file: | CytoSNP-850Kv1-2_NS550_B3_ClusterFile_GS2011.egt |
| **Genome build name: GRCh37** | Ensembl version: 74 |
| GTC file: | 204556110015_R04C01.gtc |
| Algorithm: | BeadArray v2 - Standard |
| Smoothing: | Backbone = 9 |
| **CGH Reporting:** | Minimum Del and Dup Size = 400 Kb<br>Minimum LOH Region Size (Mb) = 5.0 |
| **Significant Clones:** | CGH Region = 10 LOH Region = 500 |
| **QC Measures:** | Pass |
| Median Log R Deviation: Ratio Intensity Signal (<0.2) | 0.13 |
| Median Call Rate: genotype calling performance estimate (>0.98 ) | 1 |
| Sample sex | Female |

| ISCN | Type | Chromosome | Start | End |
|---|---|---|---|---|
| **3p21.31-3p21.1** | **LOH** | 3 | 49,641,049 | 52,968,801 |
| Loss of heterozygosity of 3328Kb (>=1Mb) on the short arm of chromosome 3. | | | | |
| **7q11.21-7q11.21** | **GAIN** | 7 | 62,047,108 | 62,705,018 |
| Copy number gain of 658Kb (<1Mb) on the long arm of chromosome 7. | | | | |
| **15q11.2-15q13.2** | **LOSS** | 15 | 22,652,330 | 30,657,952 |
| Copy number loss of 8006Kb (>=1Mb) on the long arm of chromosome 15. | | | | |

The significance of the Illumina molecular karyotyping findings should be interpreted by the principle investigator for research purposes only and include consideration of cell origin, culture conditions and experimental questions. Copy number gains and losses are reported at 400 kbp size or larger.
Copy number gain on chromosome 7q and LOH on 3p are consistent with those reported in the parent line on May 18, 2018 SC_H9_CT.

| | |
|---|---|
| Array processing: | Lisa LaBelle, MS, MB (ASCP) |
| Data analysis and sign out: | Judy Brown, PhD, CG, MB (ASCP) |

Warning: Results reported herein are for research use only and not to be used for patient diagnosis or treatment.

UCONN UNIVERSITY OF CONNECTICUT — University of Connecticut Chromosome Core — Chromosome Core

**Case Report**

| | |
|---|---|
| **Sample ID:** | **CC20-12** |
| **Sample Name:** | **H9 AG #13** |
| **Sample arrival date:** | July 11, 2020 |
| **Experiment date:** | August 26, 2020 |
| **Report date:** | April 29, 2020 |
| **Microarray type:** | Illumina CytoSNP-850K v1.2 |
| **Microarray Barcode:** | 204556110015 |
| SNP manifest file: | CytoSNP-850Kv1-2_NS550_B3.bpm |
| Annotation DB: | BG_Annotation_Ens74_20180801.db |
| SNP cluster file: | CytoSNP-850Kv1-2_NS550_B3_ClusterFile_GS2011.egt |
| **Genome build name: GRCh37** | Ensembl version: 74 |
| GTC file: | 204556110015_R05C01.gtc |
| Algorithm: | BeadArray v2 - Standard |
| Smoothing: | Backbone = 9 |
| **CGH Reporting:** | Minimum Del and Dup Size = 400 Kb<br>Minimum LOH Region Size (Mb) = 5.0 |
| **Significant Clones:** | CGH Region = 10 LOH Region = 500 |
| **QC Measures:** | Pass |
| Median Log R Deviation: Ratio Intensity Signal (<0.2) | 0.12 |
| Median Call Rate: genotype calling performance estimate (>0.98 ) | 1 |
| Sample sex | Female |

| ISCN | Type | Chromosome | Start | End |
|---|---|---|---|---|
| **3p21.31-3p21.1** | **LOH** | 3 | 49,641,049 | 52,968,801 |
| Loss of heterozygosity of 3328Kb (>=1Mb) on the short arm of chromosome 3. | | | | |
| **7q11.21-7q11.21** | **GAIN** | 7 | 62,047,108 | 62,699,114 |
| Copy number gain of 652Kb (<1Mb) on the long arm of chromosome 7. | | | | |
| **15q11.2-15q13.1** | **LOSS** | 15 | 22,750,305 | 28,535,266 |
| Copy number loss of 5785Kb (>=1Mb) on the long arm of chromosome 15. | | | | |

The significance of the Illumina molecular karyotyping findings should be interpreted by the principle investigator for research purposes only and include consideration of cell origin, culture conditions and experimental questions. Copy number gains and losses are reported at 400 kbp size or larger.
Copy number gain on chromosome 7q and LOH on 3p are consistent with those reported in the parent line on May 18, 2018 SC_H9_CT.

| | |
|---|---|
| Array processing: | Lisa LaBelle, MS, MB (ASCP) |
| Data analysis and sign out: | Judy Brown, PhD, CG, MB (ASCP) |

Warning: Results reported herein are for research use only and not to be used for patient diagnosis or treatment.

UCONN UNIVERSITY OF CONNECTICUT — University of Connecticut Chromosome Core — Chromosome Core

**Case Report**

| | |
|---|---|
| **Sample ID:** | **CC20-13** |
| **Sample Name:** | **H9 PWS #2** |
| **Sample arrival date:** | July 11, 2020 |
| **Experiment date:** | August 26, 2020 |
| **Report date:** | April 29, 2020 |
| **Microarray type:** | Illumina CytoSNP-850K v1.2 |
| **Microarray Barcode:** | 204556110015 |
| SNP manifest file: | CytoSNP-850Kv1-2_NS550_B3.bpm |
| Annotation DB: | BG_Annotation_Ens74_20180801.db |
| SNP cluster file: | CytoSNP-850Kv1-2_NS550_B3_ClusterFile_GS2011.egt |
| **Genome build name: GRCh37** | Ensembl version: 74 |
| GTC file: | 204556110015_R06C01.gtc |
| Algorithm: | BeadArray v2 - Standard |
| Smoothing: | Backbone = 9 |
| **CGH Reporting:** | Minimum Del and Dup Size = 400 Kb<br>Minimum LOH Region Size (Mb) = 5.0 |
| **Significant Clones:** | CGH Region = 10 LOH Region = 500 |
| **QC Measures:** | Pass |
| Median Log R Deviation: Ratio Intensity Signal (<0.2) | 0.13 |
| Median Call Rate: genotype calling performance estimate (>0.98 ) | 1 |
| Sample sex | Female |

| ISCN | Type | Chromosome | Start | End |
|---|---|---|---|---|
| **3p21.31-3p21.1** | **LOH** | 3 | 49,641,049 | 52,968,801 |
| Loss of heterozygosity of 3328Kb (>=1Mb) on the short arm of chromosome 3. | | | | |
| **7q11.21-7q11.21** | **GAIN** | 7 | 62,047,108 | 62,699,114 |
| Copy number gain of 652Kb (<1Mb) on the long arm of chromosome 7. | | | | |
| **15q11.2-15q13.1** | **LOSS** | 15 | 23,667,412 | 28,535,266 |
| **15q13.2-15q13.3** | **LOSS** | 15 | 30,507,461 | 32,931,921 |
| Copy number loss of 4,867,855 bp (4,868 Kb) on the long arm of chromosome 15 at bands q11.2-q13.1 and a copy number loss of 2,424,461 bp (2,424 Kb) on the long arm of chromosome 15 at bands q13.2-q13.3. | | | | |

The significance of the Illumina molecular karyotyping findings should be interpreted by the principle investigator for research purposes only and include consideration of cell origin, culture conditions and experimental questions. Copy number gains and losses are reported at 400 kbp size or larger.
Copy number gain on chromosome 7q and LOH on 3p are consistent with those reported in the parent line on May 18, 2018 SC_H9_CT.

| | |
|---|---|
| Array processing: | Lisa LaBelle, MS, MB (ASCP) |
| Data analysis and sign out: | Judy Brown, PhD, CG, MB (ASCP) |

Warning: Results reported herein are for research use only and not to be used for patient diagnosis or treatment.

UConn Chromosome Core
(860) 486-3618 https://cgi.uconn.edu/

**Supplemental Figure 2.** Report from CytoSNP array. H9Δmat15q_1 is designated as H9 AS #13, H9Δmat15q_2 is designated as H9 AS #22, H9Δpat15q is designated as H9 PWS #2.

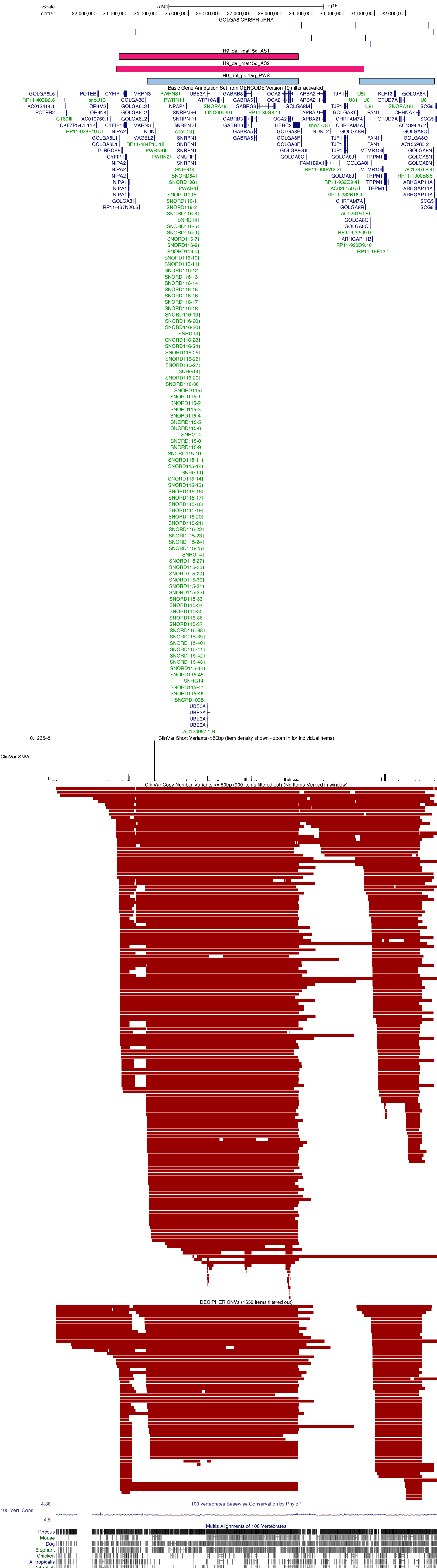


**Supplemental Figure 3.** UCSC Genome Browser shot of the chromosome 15q locus. First track displays GOLGA8 CRISPR gRNA binding sites. Second, third, and fourth tracks show deletion contained within each cell line, as determined by CytoSNP. Fifth track displays GENCODEv19 gene annotations; genes in dark blue are protein coding, genes in green are non-coding, arrows indicate direction of gene transcription. Sixth track displays short (<50 bp) variants found in the ClinVar database as a density plot. Seventh track displays ClinVar copy number variants (CNVs); for clarity, only deletions with pathogenic or likely pathogenic annotations are shown. Eighth track shows DECIPHER CNVs, only deletions with pathogenic or likely pathogenic annotations are shown. Ninth track shows conservation scores. Tenth track shows alignment of sequences to other vertebrates.

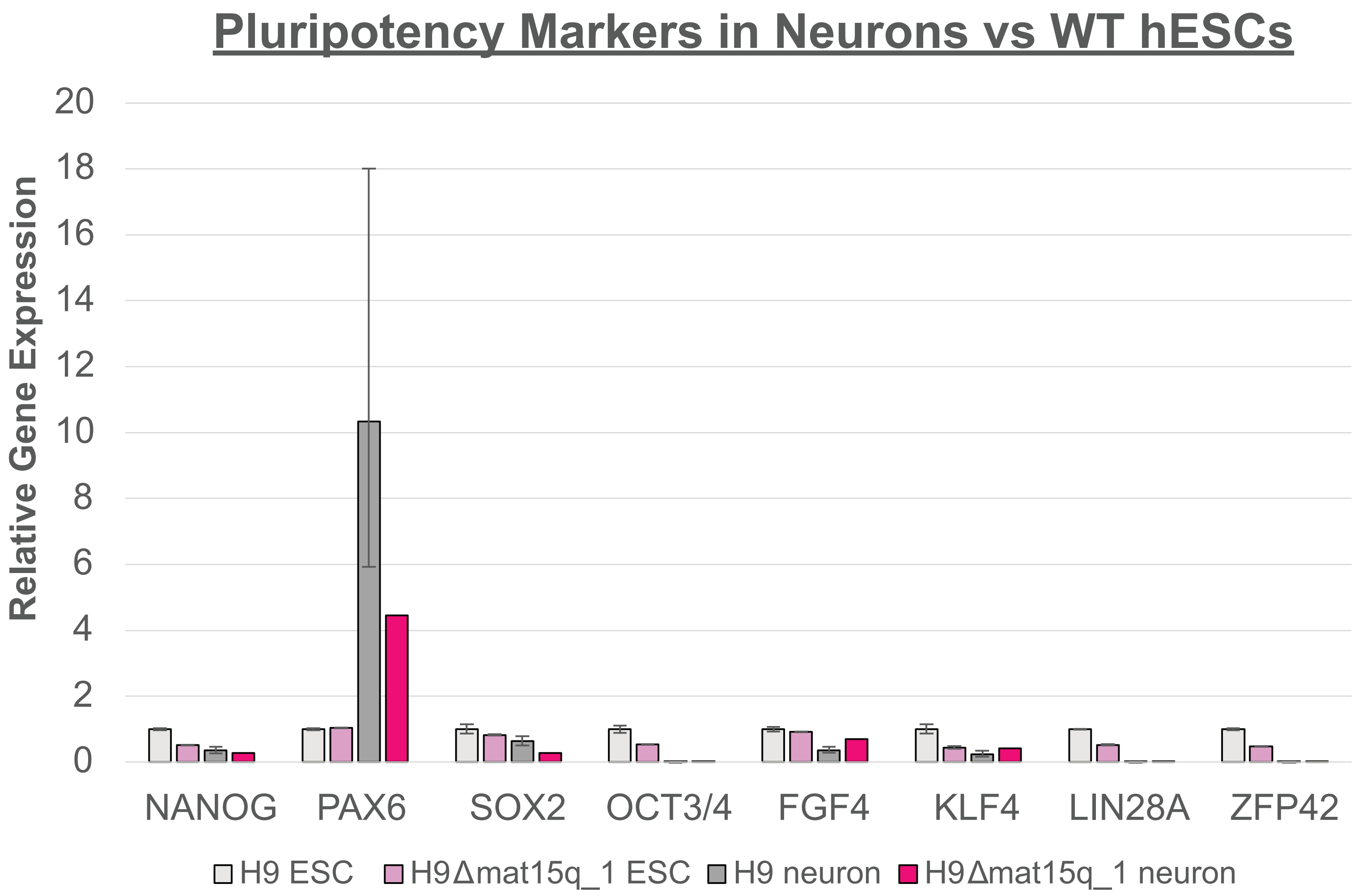


**Supplemental Figure 4.** qPCR analysis of pluripotency markers in maternal deletion line as mature 10-week neurons (n = 1-2 biological replicates). RNA expression is presented relative to the parental H9 line as ESCs. Error bars represent standard error of the mean ΔCt.
